# Rewinding the tape: historical contingency and functional constraints have shaped the evolution of APikL virulence effectors in the blast fungus

**DOI:** 10.1101/2025.08.12.669671

**Authors:** Thorsten Langner, Abbas Maqbool, Sophien Kamoun

**Affiliations:** Max-Planck-Institute for Biology, 72076, Tübingen, Germany; The Sainsbury Laboratory, University of East Anglia, Norwich Research Park, Norwich, UK; Biophysical Analysis Platform, John Innes Centre, Norwich Research Park, Norwich, NR4 7UH, UK

## Abstract

Protein evolution is influenced by historical contingencies and functional constraints, but their combined impact on rapidly diversifying pathogen virulence effectors remains poorly understood. Here, we combined ancestral state reconstructions and functional assays to recapitulate the evolution of the MAX-fold effector protein APikL2 of the plant pathogenic blast fungus *Magnaporthe* (syn. *Pyricularia*) *oryzae*, focusing on the ancestral and functionally critical amino acid residue D66 (Asp, Codon: GAT). “Rewinding the tape” experiments based on ancestral sequence resurrection revealed that, out of the seven potential amino acid substitutions derived from single nucleotide polymorphisms, only the naturally occurring D66N (Asp to Asn, GAT to AAT) expanded the binding spectrum to host plant proteins of the heavy metal associated (HMA) family. In contrast, three of the nonsynonymous substitutions were deleterious resulting in loss of binding to HMA proteins. Additionally, we identified three cases of homoplasy in the APikL effector family, involving HMA-binding interfaces, indicating recurrent convergent evolution. Our findings suggest an experimental framework for predicting evolutionary outcomes of pathogen effector—host target interactions with implications for plant disease resistance breeding.

## Introduction

Plant pathogens are engaged in persistent coevolutionary conflict with their hosts. In recent years, comparative genomics across multiple timescales have provided invaluable insights into patterns of pathogen adaptive evolution, including numerous cases in the large repertoires of effector proteins that are secreted by plant pathogens to mediate virulence (Latorre et al., 2020; Le Naour-Vernet et al., 2023; Kim et al., 2019; Yan et al., 2023; Yoshida et al., 2016). Typically, signatures of adaptive evolution in plant pathogen effectors are driven by either evasion of host immunity or adaptation to new host targets (Kanzaki et al., 2012; Li et al., 2023; Zess et al., 2022; Lin et al., 2020; Dong et al., 2014). However, the constraints that restrict adaptive evolution of plant pathogens and their virulence effectors remain poorly understood despite their implications for disease management. In this paper, we address this understudied question: what are the contingencies and functional constraints that determine the evolution of plant pathogen virulence effectors? Addressing this question should prove valuable for evaluating the diversification, predicting future evolutionary paths, and informing surveillance of emerging plant pathogens. Ultimately, such knowledge could help to guide pathogen surveillance and disease resistance development in crops.

In the blast fungus, *Magnaporthe* (syn. *Pyricularia*) *oryzae*, effector gene evolution is a hallmark of host adaptation and specialization. This multihost pathogen consists of an assemblage of genetically differentiated lineages that tend to be associated with particular host genera covering more than 50 species of grasses. Despite their overall genetic conservation in *M. oryzae* as a species, effector genes tend to display lineage-specific presence-absence patterns and pervasive diversification that resulted in dynamic effector families (Kim et al., 2019; De La Concepcion et al., 2024; Khang et al., 2008; Were et al., 2025; Zdrzałek et al., 2024). One example is the AVR-Pik like (APikL) family, consisting of six-genes and named after the well-studied rice blast fungus effector AVR-Pik (Bentham et al., 2021; Yoshida et al., 2009, Kanzaki et al., 2012). APikL effectors share a common core structure known as the MAX fold (Magnaporthe AVRs and ToxB) (de Guillen et al., 2015). Many of the functionally characterized MAX effectors, such as AVR-Pik, APikL2, AVR1-CO39, AVR-Pia and PWL2 bind plant HMA (heavy metal associated) domain containing proteins to promote infection of the host plant (Maidment et al., 2021; Bentham et al., 2021; Cesari et al., 2013; Guo et al., 2018; Were et al., 2025; Zdrzałek et al., 2024).

APikL effectors (AVR-Pik, AVR-PikL1-5) are typically associated with particular host-adapted lineages of *M. oryzae* (Bentham et al., 2021). The exception is APikL2, a highly conserved effector that is present across all blast fungus lineages. Previously, we investigated the biochemical and structural basis of adaptive evolution of APikL2 in a global population dataset of blast fungus lineages from 13 grass host species. This revealed that a single amino acid polymorphism D66N (Asp to Asn, GAT to AAT) expanded the binding spectrum of APikL2 to HMA domain containing host proteins by modulating the hydrogen bonding network in one of the HMA-binding interfaces of APikL2. We thus provided a biophysical mechanism for effector adaptation to host susceptibility proteins in the APikL family (Bentham et al., 2021).

Here, we experimentally challenged our conclusions by applying ancestral state reconstruction to “rewind the tape” and functionally test the evolutionary outcomes that could have arisen from single nucleotide mutations of the ancestral codon encoding Aspartate 66 (GAT; D66). Remarkably, of the seven possible non-synonymous mutations, only D66N (GAT→ AAT) expanded the binding spectrum of APikL2 to HMA domain proteins. We conclude that evolution of position 66 in APikL2 is constrained by the identity of the ancestral codon and the limited ability of amino acid variants at position 66 to confer binding to host-target proteins while maintaining structural integrity of the protein. In addition, we noted three cases of homoplasy in the APikL effector family indicating parallel evolution and functionally constrained diversification of key amino acids at effector-host target binding interfaces. Our findings provide a mechanistic evolutionary framework for recapitulating pathogen effector evolution. We propose that such a framework can inform the prediction of possible evolutionary outcomes for pathogen effector-host target interactions with important consequences for pathogen-guided breeding for disease resistance.

## Results

### Only the naturally occurring D66N amino acid substitution leads to an expansion of host-target binding in ancAPikL2

To test the possible evolutionary outcomes that could have contributed to the diversification of APikL2 in position 66, we reconstructed and synthesized (i.e., resurrected) the ancestral variant of APikL2 that precedes allelic diversification (ancAPikL2; Fig. 1A). Of the seven polymorphic residues that differentiate modern APikL2 effectors, the ancestral variant contains Ile26, Val63, Asp66, Asp77, Gly107, Ile108, and Pro111 (Fig. 1A). Four of these residues diverge between the previously tested variants, ApikL2A and APikL2F (Fig. 1B). In this ancestral sequence context, we introduced every non-synonymous, single nucleotide mutation in the codon encoding D66 to reconstruct the possible evolutionary outcomes. This resulted in seven APikL2 variants with amino acid substitutions from Asp to Ala, Tyr, His, Glu, Val, Gly, and Asn, with only the latter observed in natural populations of the blast fungus (Fig. 1). We then tested the effect of these mutations on host-target binding in a comparative pairwise yeast two-hybrid assay using the previously identified host target HMA domain sHMA94 (Bentham et al., 2021). As a control, we included sHMA25 that binds to both modern variants, APikL2A and APikL2F. Three mutations (D66 to Y66, H66 or V66) were deleterious and disrupted binding to sHMA25 while another three mutations (D66 to A66, E66 or G66) had only minimal effects on the interaction with the host target control. Intriguingly, only the D66N mutation led to a gain of binding phenotype, and thus an expansion of the host target binding range in ancAPikL2 (Fig. 1B).

**Fig. 1.**
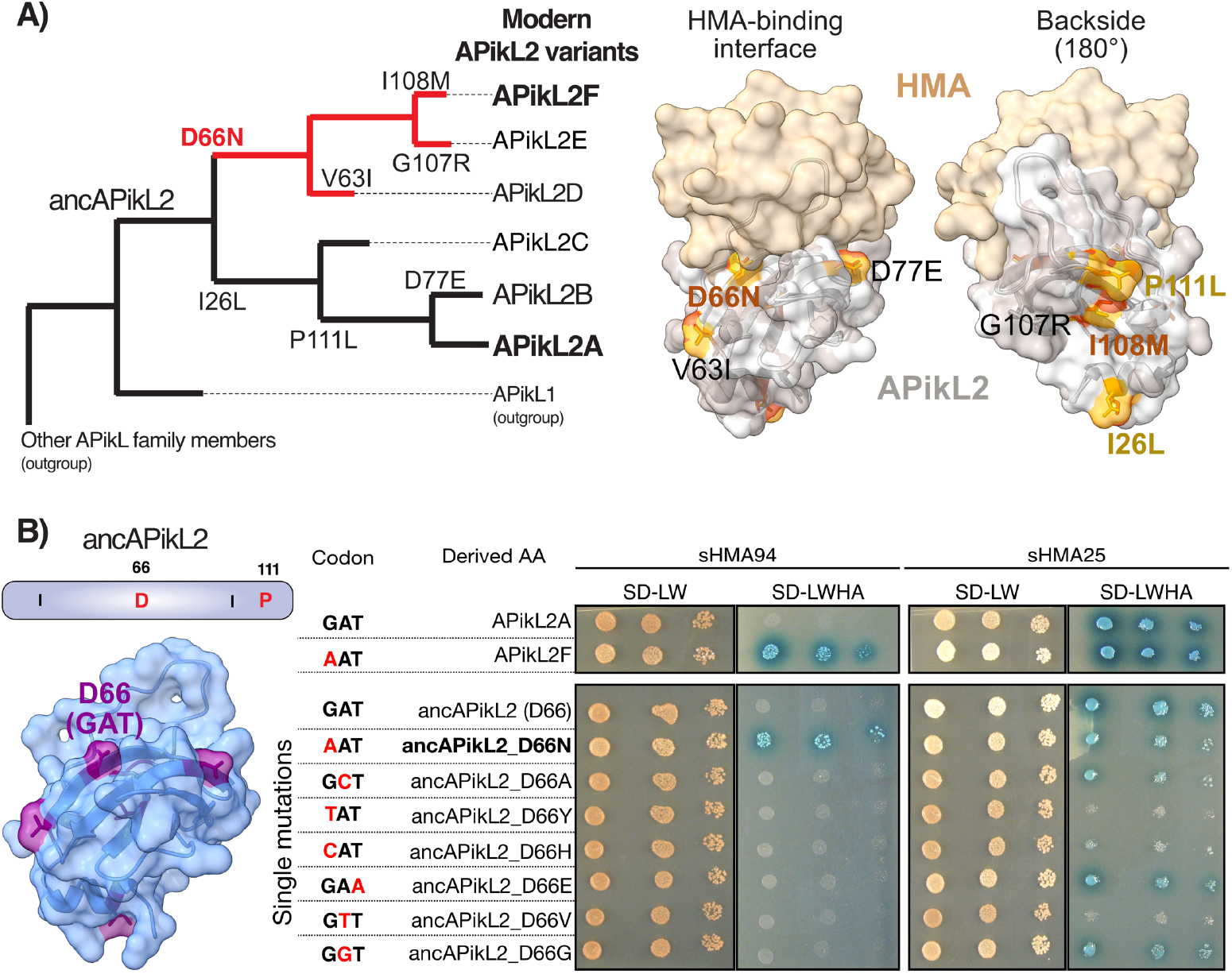
Only the D66N substitution leads to a target range expansion in ancAPikL2. **A)** Seven non-synonymous mutations contribute to the differentiation of modern APikL2 variants from the ancestral APikL2 state. Left: Position of the ancAPikL2 variant in the tree and derived mutations in modern APikL2 variants. Right: Position of polymorphic amino acids relative to the HMA-effector binding interface. Amino acid positions that led to the differentiation of APikL2A and APikL2F are shown in brown and beige, respectively. Other polymorphic residues are shown in black. **B)** Reconstruction of possible evolutionary outcomes by single nucleotide mutations at the codon encoding D66 (GAT). Mutated nucleotides are indicated in red and the corresponding amino acid substitution in ancAPikL2 shown in the center. Yeast two-hybrid interaction screen of ancAPikL2 variants with the HMA domains sHMA94 and the control sHMA25 (right). For each drop, 4 *µ*l of a serial dilution of OD600= 1, 0.1, and 0.01 were applied (from left to right).

### Homoplasy at key amino acid residues at the HMA binding interface indicate recurrent convergent evolution in the APikL effector family

Following the experimental evidence for functional constraints that determine the evolutionary outcome of APikL2 D66, we extended our analysis to the entire APikL effector family. We annotated every amino acid position that contributes to allelic diversification in the APikL family and extracted corresponding codons to identify substitutions that may diversify under evolutionary constraints. In addition to the previously identified D66N substitution, we identified three cases of homoplasy (i.e., parallel evolution) leading to the independent acquisition of P47A, V63I, and M78L mutations (Fig 2A and C). Intriguingly, these amino acid substitutions were caused by either the same mutation (indicated in blue) or independent mutations (red) leading to the same amino acid. In position 47, AVR-Pik and APikL4 acquired the P47A by a single mutation from CCT (P) to GCT (A). Position 78 mutated from an ancestral ATG (M) codon to either CTG (APikL1) or TTG (APikL4 and APikL5), both leading to the acquisition of M78L. In addition, position 63 acquired at least two mutations leading to the substitution of GTA (V) to ATT (I; ApikL2) and ATC (I; APikL1). Two of the identified residues, position 47 and 78, are located at the HMA-target binding interface (Fig 3B) suggesting that convergent evolution of these amino acids may be driven by functional constraints imposed by their HMA-host targets.

**Fig. 2.**
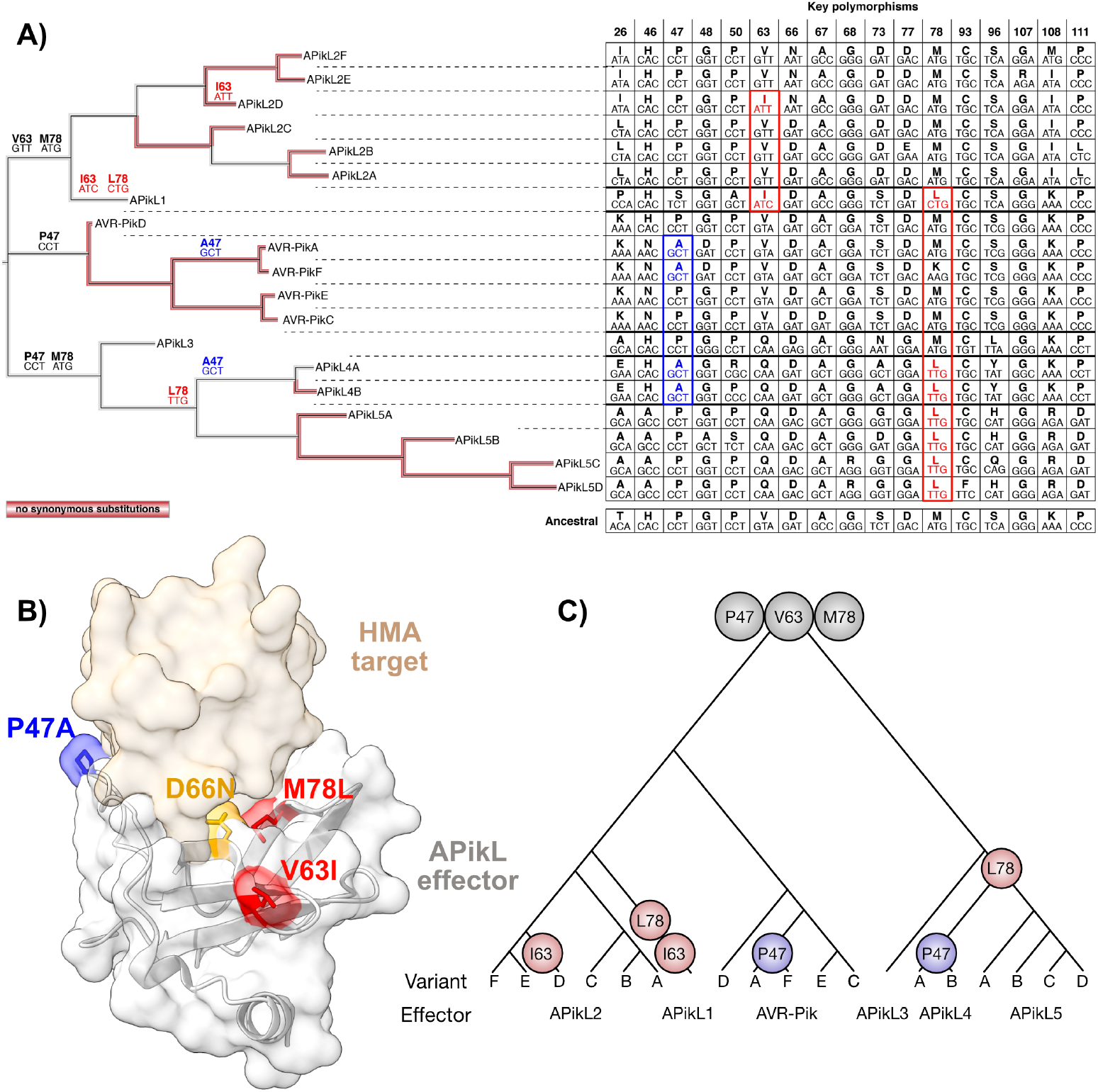
Homoplasy suggests functionally constrained diversification of key amino acids in the APikL effector family. **A)** Ancestral sequence reconstruction of codons that encode diversifying amino acids in the APikL effector family. Mutations that lead to independent acquisition of homoplasic amino acid substitutions are highlighted in blue (caused by the same nucleotide mutation) and red (caused by different nucleotide mutations leading to the same amino acid). The ancestral state is indicated in black in the tree and at the bottom of the codon table. **B)** Position of amino acid residues under convergent evolution highlighted in the crystal structure of APikL2A in complex with sHMA25 (PDB accession: 7NLJ). **C)** Schematic representation of homoplasy in the APikL effector family.

## Discussion

Plant pathogen virulence effector proteins are in coevolutionary conflict with host-target proteins leading to rapid adaptive evolution (Bentham et al., 2021; Maidment et al., 2021; Were et al., 2025; Dong et al., 2014). While these processes are extensively studied from either an evolutionary or mechanistic perspective, current efforts aim to combine these approaches to develop integrated mechanistic evolutionary frameworks to understand and predict evolutionary outcomes (Schornack and Kamoun, 2023).

Here, we describe constrained evolution of key amino acids in a conserved effector-host target binding interface of the multihost pathogen *M. oryzae*. By leveraging ancestral sequence reconstruction and resurrection of ancestral effector variants we demonstrate experimentally that functional constraints imposed by the host target limit the diversification of a key amino acid in the host-target binding interface of the effector APikL2, confirming previous observations obtained from population genomics data of field isolates (Bentham et al., 2021). In addition, we detected three cases of homoplasy across the entire APikL effector family indicative of convergent evolution. Together, our data suggest that evolutionary constraints have shaped the diversification of effector-host target binding interfaces.

Multihost pathogens frequently undergo host jumps or host range expansion and are thus exposed to changing host environments (Thines, 2019; Inoue et al., 2017; Ploch et al., 2022). As a consequence, pathogen virulence effectors constantly need to adapt to a diverse host immune systems but also to diversifying effector target families (Kanzaki et al., 2012; Bentham et al., 2021; Sacristán et al., 2021; Le Naour-Vernet et al., 2023; Maidment et al., 2021). While the interplay of effectors with plant immune receptors is relatively well-studied, our understanding of evolutionary constraints and selection pressure imposed by effector virulence targets is lacking behind. Reasons for this lack of understanding include i) that virulence outcomes of an interaction of single effector molecules with their targets are often quantitative and therefore harder to detect than qualitative activation of immune responses (avirulence) and ii) because classical selection tests (e.g., dN/dS ratios) can fail because of heavily skewed mutation ratios (e.g., when an effector accumulates non-synonymous mutations in absence of synonymous ones). Therefore, experimental validation in combination with alternative predictive methods are needed to understand adaptive pathogen effector evolution.

Parallel and convergent evolution are often seen as the most compelling evidence for evolutionary outcomes driven by natural selection as both of these processes lead to the independent emergence of adaptive traits – an outcome that is unlikely to happen by chance (Cerca, 2023). Previous studies of MAX-fold virulence effectors of the blast fungus provided compelling evidence for functional convergence (De Guillen et al., 2015; Guo et al., 2018; Maqbool et al., 2015; Ortiz et al., 2017; De La Concepcion et al., 2018). These effector proteins are sequence unrelated, but share a common core structure. In addition, several of these structurally related effectors converged to bind heavy metal associated (HMA) domains of host-target proteins, indicating convergent evolution of host-target binding (Bentham et al., 2021; Cesari et al., 2014; Were et al., 2025; Zdrzałek et al., 2024).

In addition, homoplasy, i.e., the parallel emergence of a specific trait that is not explained by a common ancestor, is a strong indicator of evolutionary constraints leading to independent acquisition of adaptive traits (Edwards et al., 2021). Here, we observed three sites within the APikL effector family for which an independent acquisition of specific amino acids indicates homoplasies. Notably, two of these amino acids (position 47 and 78) and the experimentally confirmed D66N mutation locate at the effector-HMA interaction interface and have been shown to be involved in host-target binding specificity (Maidment et al., 2021; Bentham et al., 2021), further pointing towards convergent evolution driven by a common host-target binding mechanism.

In conclusion, our study provides experimental and evolutionary evidence for functional constraints that limit the diversification of pathogen effector interaction interfaces. Rewinding the tape using ancestral state reconstruction and gene synthesis provides an opportunity to test possible evolutionary outcomes and to develop a mechanistic evolutionary framework for recapitulating evolution, enabling the development of predictive models for the diversification of pathogen effector proteins. Such models could be used in the future to predict the emergence of virulence traits in pathogen populations and can guide the development of disease management strategies, such as the predictive bioengineering of plant immune receptors.

## Material and Methods

### Reconstruction of ancestral states in the APikL effector family and resurrection of ancAPikL2

We extracted genomic sequences of all APikL effector family members from 107 *M. oryzae* genomes representing 10 diverse genetic lineages (as described in Bentham et al., 2021) and generated multiple sequence alignments using MUSCLE (Edgar, 2004). We then generated a codon-based, maximum likelihood tree with 1000 bootstrap iterations of the family using MEGA 7 (Kumar et al., 2016). Next, we performed a phylogenetic reconstruction and determined ancestral states using a codon based maximum likelihood analysis with codeml (included in the PAML v4.9 package) according to (Yang et al., 1997 and Yang, 2007). Ancestral sequences of internal nodes involved in allelic diversification of the APikL effector family as well as ancestral codons in polymorphic sites were extracted from codeml. Ancestral sequences of APikL2 were synthesized by Genewiz from Azenta Life Sciences and cloned into the Matchmaker 3 pGADT7 yeast two-hybrid plasmids (Takara Bio Inc.). We next introduced every possible non-synonymous single nucleotide substitution in the ancestral state predating the divergence of APikL2 alleles (Fig 1A and 2) to generate possible evolutionary outcomes in the codon encoding for Asp in position 66.

### Modelling of the ancAPikL2 structure

The structure of ancAPikL2 was predicted by Alphafold3 (Abramson et al., 2024) using default parameters of the Alphafold 3 server (https://alphafoldserver.com/). We confirmed the models by superimposition of the predicted structures with experimental structures determined for the modern APikL2 variants APikL2A (PDB: 7NLJ) and APikL2F (PDB: 7NMM) in complex with host target heavy metal associated (HMA) domains in ChimeraX (Meng et al., 2023; Pettersen et al., 2021; Goddard et al., 2018). Visualization of protein structures and derived mutations in APikL2 was also done in ChimeraX. Experimental structures of APikL2F and APikL2A in complex with the HMA domain HMA94 and HMA25, respectively (Bentham et al., 2021), were used to visualize derived mutations in modern APikL2 alleles.

### Interaction of ancAPikL2 variants with HMA domains HMA94 and HMA25

All synthesized ancAPikL2 variants were cloned into the pGADT7 yeast two-hybrid plasmids (Takara Bio Inc.). Host (3,10–13)target HMA domains were cloned into pGBKT7 (Takara Bio Inc.) as described previously (Bentham et al., 2021; Langner et al., 2021). Interaction of effector variants and HMA domains was tested using the Matchmaker 3 yeast two-hybrid kit (Takara Bio Inc.) following the manufacturer’s instructions. Each individual effector-HMA combination was co-transformed into competent *Saccharomyces cerevisiae* Y2H Gold cells (Takara Bio USA) and selected on SD^-Leu-Trp^ (synthetic dropout) medium. Single colonies were selected and grown overnight at 28°C and 200 rpm to an OD_600_ of 1–2 in liquid SD^-Leu-Trp^ medium. Cultures were then used to prepare serial dilutions of OD600 1, 1^−1^, 1^−2^. Of each dilution 4 μl were replica plated on SD^-Leu-Trp^ and SD^-Leu-Trp-Ade-His^ containing the chromogenic substrate X-α-Gal. Pictures were taken after 96 h incubation at 28°C. HMA25 was used as a positive control that binds to both modern APikL2 variants, APikL2A and ApikL2F.

### Detection of homoplasy in the APikL effector family

The maximum likelihood tree generated for ancestral state reconstructions was used to visualize derived mutations across the APikL effector family. The red color in the tree indicates branches where no synonymous mutations occur (reprinted from Bentham et al., 2021). Branches where mutations occur that lead to independent emergence of amino acid polymorphisms are indicated in blue (same nucleotide mutation leads to the same derived amino acid) and red (different nucleotide substitutions lead to the same derived amino acid). Nucleotide sequences of all APikL-family members and ancestral states were translated into amino acid sequences and multiple sequence alignments were generated using Clustal Omega (Sievers and Higgins, 2014). Codon and amino acid positions where homoplasy was detected are indicated in blue and red.

## Acknowledgements

We thank all members of the Kamoun group, especially Andres Posbeyikian, the BLASTOFF team at the Sainsbury Laboratory, and Hernan Burbano and Sergio Latorre at University College London for valuable discussions.

## Author contribution

**Conceptualization:** SK, TL.

**Formal analysis:** AM, TL.

**Visualization:** TL.

**Supervision:** TL, SK.

**Writing – original draft:** TL

**Writing – review & editing:** SK, TL, AM.

**Project administration:** TL, SK.

**Funding acquisition:** SK, TL.

